# Insulin Prevents Fat Loss and Promotes Muscle Loss During Intermittent Fasting in Obesity

**DOI:** 10.1101/2025.06.21.660875

**Authors:** Daniel M. Marko, Meghan O. Conn, Joseph F. Cavallari, Helena Neudorf, Alexandra P. Steele, Brittany M. Duggan, Breanne T. McAlpin, Nicole G. Barra, Thomas J. Hawke, Jonathan P. Little, Jonathan D. Schertzer

## Abstract

Elevated blood glucose, insulin, and insulin resistance are associated with obesity and type 2 diabetes (T2D). High blood insulin levels blunt lipolysis and promote lipogenesis, and thus weight gain. Intermittent fasting (IF) has emerged as a weight loss strategy that also lowers blood glucose and improves insulin resistance. However, some people with obesity, or T2D have less fat loss and more lean mass loss after IF. It is not known why some people lose more fat or muscle during IF. We hypothesized that features of obesity, such as high insulin and insulin action in adipocytes dictate less adipose loss and more muscle loss during IF. In humans, we found that people living with obesity and higher blood insulin lost more lean mass after a 48-hour fast. Chronic elevation of insulin in obese mice lowered adipose loss and promoted muscle loss after 10 weeks of 5:2 IF in obese mice, while concurrently lowering adipose tissue interferon regulatory factor 4 (IRF4) expression. Whole-body and adipocyte-specific deletion of *Irf4* in mice phenocopied chronic hyperinsulinemia, resulting in less fat loss and greater muscle loss after 10 weeks of 5:2 IF, which occurred in mouse models of equal and reduced caloric intake during IF. Therefore, hyperinsulinemia and suppression of adipocyte IRF4 promote muscle loss over fat loss during IF.

**Significance Statement:** It is not known why some people lose muscle during IF. In humans, we found that high blood insulin during obesity correlated with increased lean mass loss after one 48-hour fast. Mechanistically, we found that hyperinsulinemia and the insulin responsive factor IRF4 within adipocytes as regulators of adipose and muscle loss during chronic IF obese mice. Therefore, insulin status and regulation of adipocyte IRF4 may be important factors to consider before prescribing IF for weight loss as lower muscle mass is known to be detrimental to metabolic and overall health status.

## Introduction

Obesity is the largest risk factor for type 2 Diabetes (T2D), which are global health problems (1). Obesity is characterized by adipose tissue expansion, where fat mass increases more than skeletal muscle mass (2). Anti-obesity drugs, such as GLP-1-receptor agonists, lower food intake, fat mass, and risk of T2D (3). Loss of skeletal muscle during medically induced weight loss is a concern, including during incretin-based drug therapy (4). Intermittent fasting (IF) is another approach to combat obesity and metabolic disease. Loss of skeletal muscle versus fat should also be considered during IF.

IF consists of many approaches involving repeated bouts of food deprivation through multiple protocols, including i) alternative day fasting (ADF), which involves fasting every other day of the week, ii) modified fasting (5:2), which limits food intake on two non-consecutive days of the week, and iii) time-restricted eating/feeding (TRE/F), which restricts calorie consumption to a certain window of time each day (5, 6). IF can decrease energy intake and increase energy expenditure in mice (7). In humans and rodents, IF can lower blood glucose, HbA1c, insulin, blood lipids and cholesterol, inflammatory cytokines, and adiposity (8–10). In people with obesity or T2D IF can lower adiposity, blood glucose, and insulin (11–14). However, IF has been reported to lower lean (or fat-free) mass in people with obesity or T2D (11, 12, 15–19). For example, TRE (with an 8-h daily feeding window) for 12 weeks did not alter fat mass but increased the loss of appendicular lean mass in people with obesity (11). Additionally, TRE for 6-18 months promoted more lean mass loss compared to caloric restriction in people with obesity and at risk for T2D (12). Taken together, these studies indicate that IF may affect the balance of fat and lean mass in people with obesity and/or T2D differently. Hence, we tested if people with obesity and elevated blood insulin lost more lean mass than fat mass during an acute fast. We found that people with obesity and higher blood insulin lost more lean mass during a 48-hour fast compared to people who were not obese. Hence, we sought to determine if hyperinsulinemia was sufficient to promote lean mass loss during IF and determine the mechanisms of action balancing muscle versus fat mass loss during IF.

High insulin levels inhibit lipolysis, by inhibiting lipases and transcriptional control of lipolysis by lowering interferon regulatory factor 4 (IRF4) in adipocytes. IRF4 is higher during acute fasting and activates lipolysis (22–25). We assessed the impact of hyperinsulinemia, and adipocyte IRF4 on fat loss versus muscle loss during IF. Our data show that repeated IF reduces body mass during high-fat diet (HFD) feeding in mice. We used different mouse models to show that IF caused weight loss when mice consumed fewer calories and when mice consumed the same number of calories during IF. Chronically high insulin and deletion of *Irf4* tipped the balance toward more muscle loss rather than fat loss during IF in obese mice.

## Results

### High insulin promotes lean mass loss during acute fasting in humans with obesity

Women and men with obesity were compared to people without obesity after they consumed a standardized mixed macronutrient breakfast drink to equilibrate baseline feeding measures and then underwent a 48-hour water-only fast (Figure 1A). People with obesity had higher serum insulin during feeding and after 48 hours of fasting (Figure 1B, C). Serum non-esterified fatty acids (NEFA) concentrations increased after 48 hours of fasting (Figure 1D). People with obesity had higher serum L-amino acids, but lower serum alanine levels in the fed and fasted state (Figure 1D-E). BMI, lean mass, and fat mass were measured in the fed state and after 48 hours of fasting in each person and change during the 48 fast was determined. People with obesity lost more body mass during fasting based on BMI (Figure 1G). People with obesity lost more lean mass, but not fat mass, during fasting (Figure 1H-I). Overall, these human study findings show that obesity and/or high insulin promotes more lean mass loss during a single bout of fasting.

**Figure 1:**
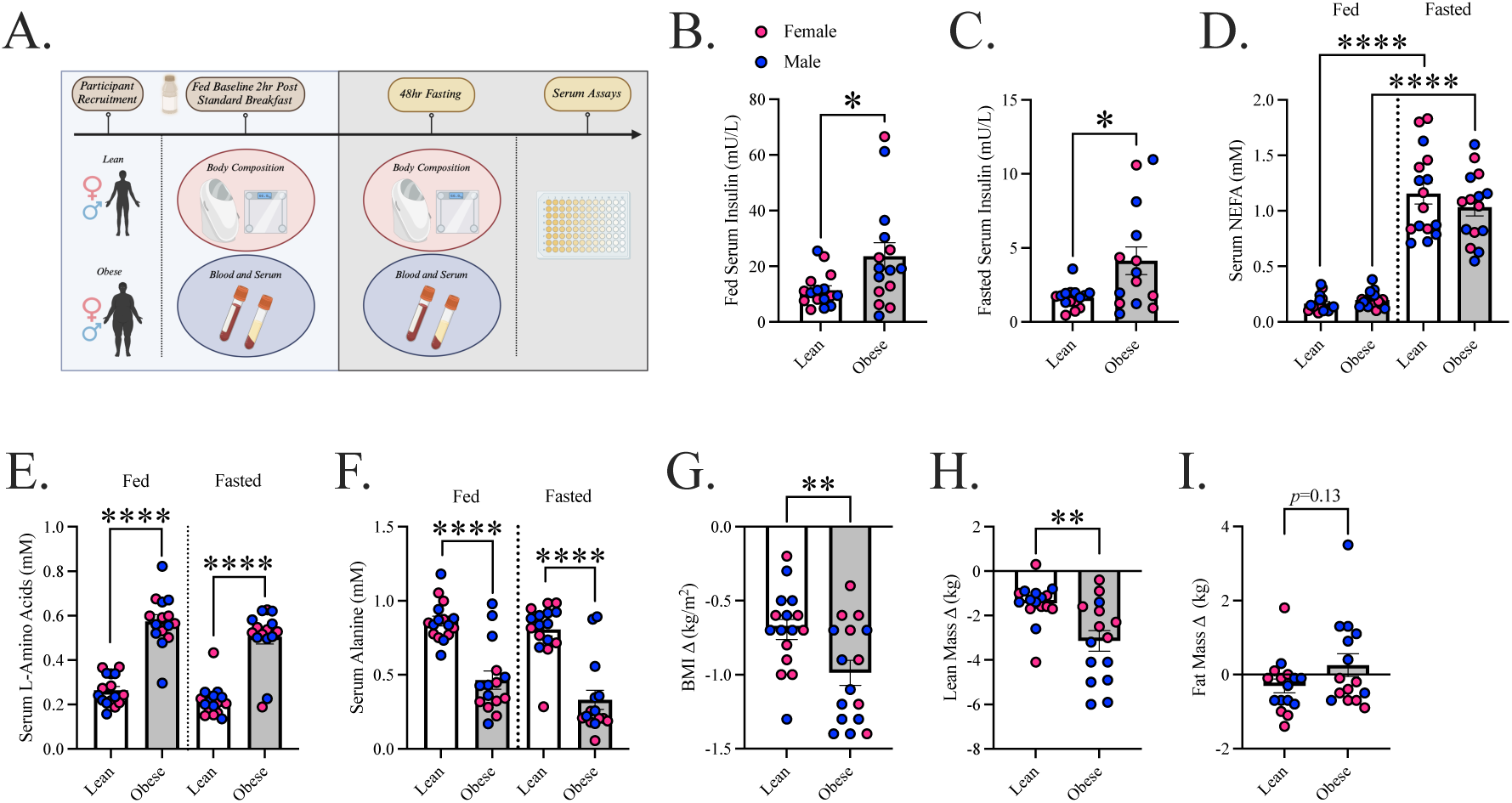
Obesity and/or High Insulin Suppress Fat Loss and Drive Lean Mass Loss During Acute Fasting in Humans. Delta values were calculated by subtracting body composition measures after 48hrs of fasting from the baseline fed measurements in each individual. A) 48hr acute fasting protocol. B) Fed baseline serum insulin. C) 48hr fasted serum insulin. D) Fed and fasted serum NEFA. E) Fed and fasted serum L-amino acids. F) Fed and fasted serum alanine. G) Delta BMI mass loss. H) Delta lean mass loss. I) Delta fat mass. Data presented as mean ± SEM (n=16). Statistical significance was determined using t-test analysis (B, C, G-I). Significance was determined using a one-way ANOVA (D, E, F). *p≤0.05, **p≤0.01, and ****p≤0.0001.

### Chronic hyperinsulinemia decreases fat loss and increases muscle loss during IF in a mouse model of obesity

Next, we built a mouse model of chronic hyperinsulinemia using subcutaneous slow-dissolving insulin pellets (INS) or sham surgery (SHAM). We focused on male mice since there were no apparent sex differences in the data from human fasting (Figure 1). INS lowered blood glucose and serum NEFA during 12 and 24 hours of fasting in mice fed a control diet (Figure 2A, B). INS lowered serum NEFA and glycerol at 20 and 60 minutes after injection of CL-316, 243, a β3-adrenergic receptor agonist, in mice fed a control diet (Figure 2D, E). These data confirmed that we had built a model of hyperinsulinemia that lowered blood glucose and blunted lipolysis. We next placed INS or SHAM mice on an obesogenic 45% HFD, where mice were fed ad-libitum (AL) or underwent 5:2 IF for 10 weeks and housed at thermoneutrality (29°C) (Figure 2F). Each 24-hour fast was water-only and occurred on 2 non-consecutive days of the week (i.e., Monday and Thursday). 5:2 IF reduced body mass compared to AL in SHAM and INS mice (Figure 2G, H). Compared to SHAM, INS mice had lower blood glucose every week when assessed after 24 hours of fasting during the IF, and the insulin pellets were replaced at week 6 (Figure 2I). Compared to SHAM, INS mice also had consistently lower random-fed blood glucose (Figure S1). INS mice with IF had significantly lower average food intake (g/mouse/day) compared to INS AL and SHAM IF (Figure 2J). IF mice had higher cage activity than AL mice (Figure 2K). To test the effect of 10 weeks of IF, values obtained after IF were subtracted from the average AL values, and tissue weights were normalized to body mass. After IF, INS mice lost less body mass than SHAM mice (Figure 2L). INS mice lost less liver mass compared to SHAM mice after IF (Figure 2M). IF lowered liver fat and triglycerides (Figure S1).

**Figure 2:**
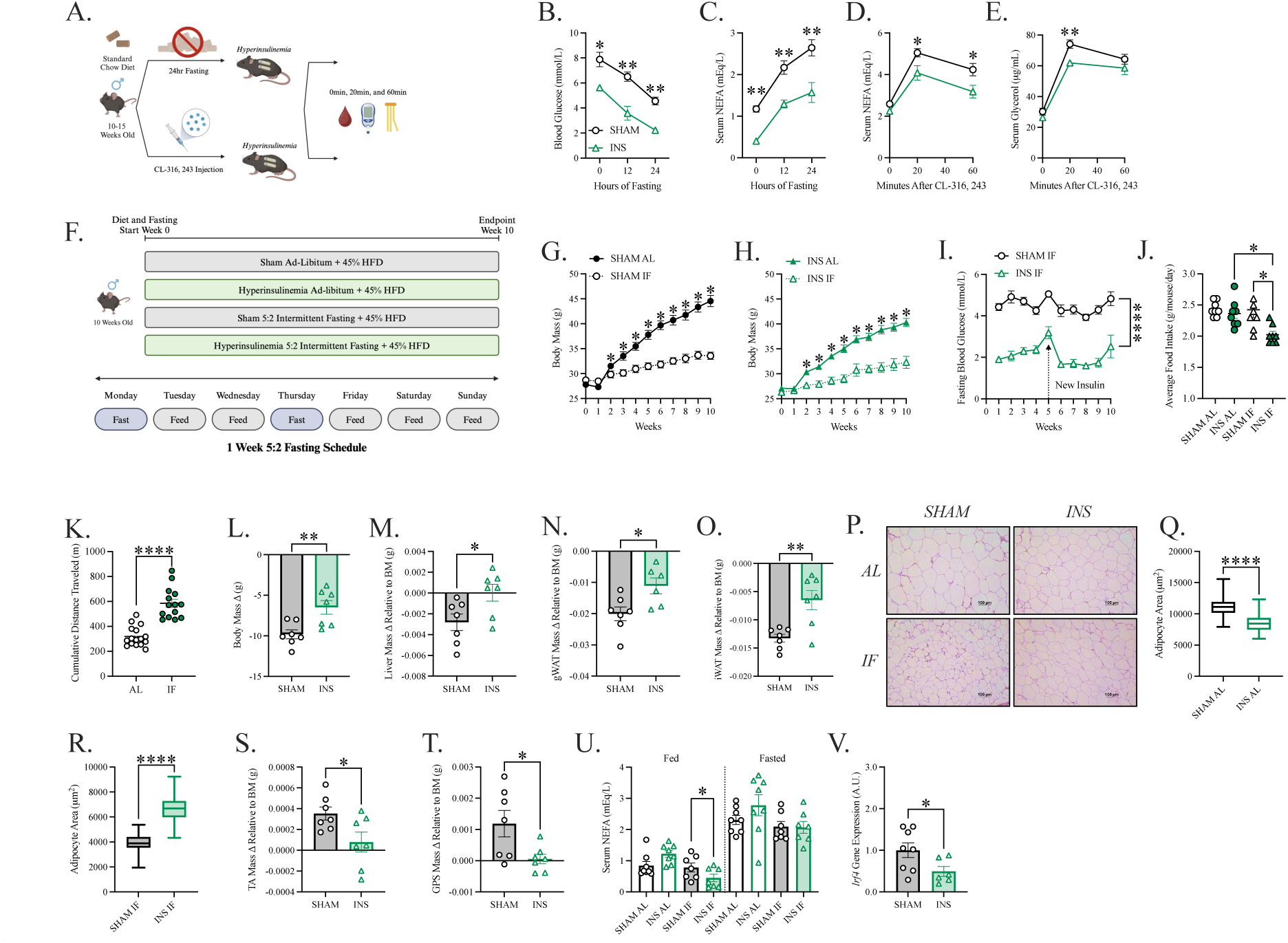
Hyperinsulinemia Blunts Lipolysis and Inhibits Fat Loss During Repeated Intermittent Fasting in Mice. All acute fasting and adrenergic experiments were done at room temperature (23°C) and mice were fed a chow diet. All 45% high-fat diet (HFD) repeated IF experiments were done at thermoneutrality (29°C). SHAM: control, INS: hyperinsulinemia, AL: ad-libitum, IF: intermittent fasting. Tissue weights were made relative to body mass and the fasting values were subtracted from the ad-libitum average of the corresponding group. A) Acute fasting and adrenergic stimulation of lipolysis protocol. B) Blood glucose levels after acute fasting. C) Serum NEFA levels after acute fasting. D) Serum NEFA after CL-316, 243 injection (i.p.). E) Serum glycerol after CL-316, 243 injection. F) 10wk 5:2 repeated intermittent fasting protocol. G) SHAM AL and IF body mass. H) INS AL and IF body mass. I) 24hr fasted blood glucose. J) Average daily food intake. K) Cage activity over 24hrs. L) Body mass delta. M) Liver mass delta. N) Gonadal white adipose tissue (gWAT) delta. O) Inguinal white adipose tissue (iWAT) mass delta. P) Adipocyte images taken at 10X objective where the scale bar represents 100mm. Q) Adipocyte cross-sectional area for AL mice. R) Adipocyte cross-sectional area for IF mice. S) Tibialis anterior (TA) muscle mass delta. T) Gastrocnemius plantaris soleus (GPS) muscle mass delta. U) Fed and 24hr fasted serum NEFA. V) gWAT *Irf4* gene expression made relative to the *Rplp0* housekeeper gene. Data presented as mean ± SEM (n=6-8). Statistical significance was determined using t-test analysis (B-I and K-V). Significance was determined with a one-way ANOVA (J). *p≤0.05, **p≤0.01, and ****p≤0.0001.

After 10 weeks of IF, INS mice lost less gonadal and inguinal white adipose tissue (gWAT, iWAT) than SHAM mice. INS mice also had larger adipocyte cross-sectional area compared to SHAM mice after IF (Figure 2N-R). Conversely, INS mice had smaller hindlimb tibialis anterior (TA) and gastrocnemius, plantaris, and soleus (GPS) muscles compared to SHAM mice after IF (Figure 2S, T). There was no difference in TA myofiber cross-sectional area between SHAM and INS mice (Figure S1). In the fed state, INS mice had lower serum NEFA, underscoring the ability of chronic hyperinsulinemia to blunt lipolysis (Figure 2U). Despite reductions in lipolytic markers in INS mice, there were no differences in serum alanine and serum glucagon was actually lower in INS mice after IF compared to AL-fed INS mice (Figure S1). It was known that insulin regulates aspects of lipolysis via the transcription factor *Irf4* in mice (22). INS mice had significantly lower *Irf4* expression in adipose tissue compared to SHAM mice (Figure 2V). Overall, these data show that chronic hyperinsulinemia in mice blunted lipolysis and tipped the balance toward less fat loss and more muscle loss during IF, which was associated with lower transcript levels of *Irf4* in adipose tissue.

### Deletion of adipocyte Irf4 decreased fat loss and increased muscle loss in an obese mouse model of IF with reduced caloric intake

Next, we tested mice with whole-body deletion of *Irf4*. Similar to chronic hyperinsulinemia, whole-body *Irf4* deletion blunted lipolysis, as evidenced by lower serum glycerol levels after 12 hours of fasting and 20 and 60 minutes after injection of CL-316, 243 in chow-fed mice (Figure 3A-C). We then placed *Irf4* knockout (KO), littermate heterozygous (HET), and littermate wild type (WT) mice on an obesogenic 45% HFD, where mice were fed AL or underwent 5:2 IF for 10 weeks and housed at room temperature (23°C) (Figure 3D). IF decreased body weight in WT, HET, and *Irf4* KO mice (Figure 3E-G). We found that a 5:2 IF protocol at room temperature paired with a 45% HFD caused a reduction in food intake during IF (Figure 3H). KO mice lost less body mass compared to HET and WT mice after IF (Figure S2). EchoMRI analysis revealed that HET and KO mice lost less fat mass and more lean mass compared to WT mice after 10 weeks of IF (Figure S2). *Irf4* HET and KO mice lost less gWAT mass and KO mice lost less iWAT mass, compared to WT mice after 10 weeks of IF (Figure 3I, J). In addition, KO mice had smaller TA muscles compared to WT and HET mice after 10 weeks of IF (Figure 3K). These data show that the deletion of *Irf4* decreased fat loss and increased muscle loss during IF in obese mice.

**Figure 3:**
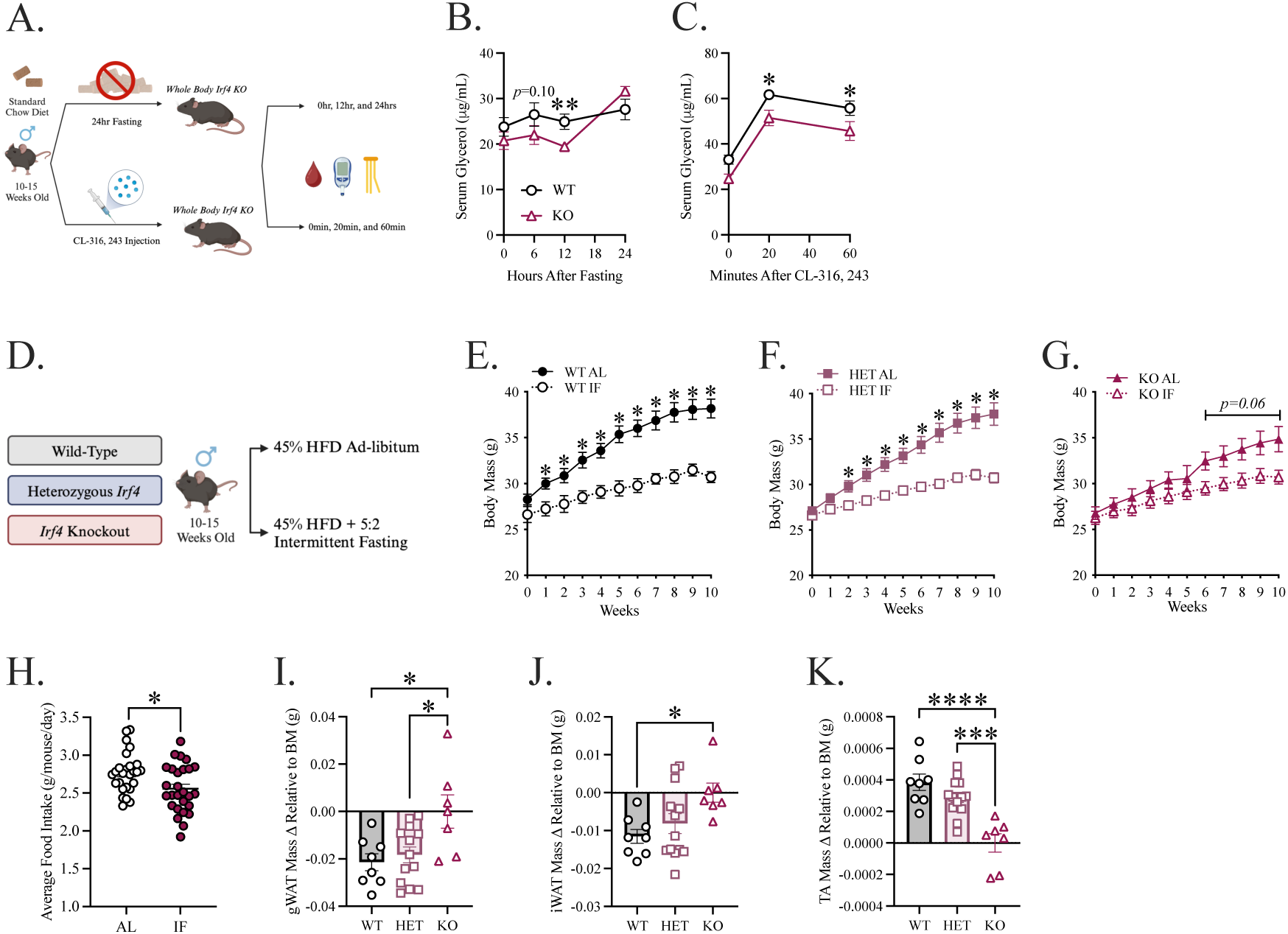
*Irf4* Impairs Lipolysis and Fat Loss While Promoting Muscle Loss in a Model of Reduced Food Intake During Intermittent Fasting. All acute fasting and adrenergic experiments were done at room temperature (23°C) and mice were fed a chow diet. All HFD repeated IF experiments were also done at room temperature (23°C). WT: wild-type control, HET: heterozygous (*Irf4^+/-^*), KO: *Irf4* knockout (*Irf4^-/-^*), and AL: ad-libitum, IF: intermittent fasting. Tissue weights were made relative to body mass and the fasting values were subtracted from the ad-libitum average of the corresponding group. A) Acute fasting and adrenergic stimulation of lipolysis protocol. B) Serum glycerol after acute 24hr fasting. C) Serum glycerol after CL-316, 243 injection (i.p.). D) 10wk 5:2 repeated intermittent fasting protocol. E) WT AL and IF body mass. F) HET AL and IF body mass. G) KO AL and IF body mass. H) AL and IF average daily food intake. I) gWAT mass delta. J) iWAT mass delta. K) TA mass delta. Data presented as mean ± SEM (n=4-25). Statistical significance was determined using t-test analysis (A-F). Significance was determined with a one-way ANOVA (G-I). *p≤0.05, **p≤0.01, ***p≤0.001, and ****p≤0.0001.

Next, we placed adipocyte-specific *Irf4* knockout mice (adipoIrf4) and littermate floxed mice (flox/flox) on an obesogenic 45% HFD, where mice were fed AL or underwent 5:2 IF for 10 weeks and housed at room temperature (23°C). IF decreased body mass in both adipoIrf4 and flox/flox mice (Figure 4A-C). Again, IF reduced daily food intake when mice were housed at room temperature (Figure 4D). AdipoIrf4 and flox/flox mice lost the same amount of body mass, but EchoMRI analyses revealed that adipoIrf4 mice lost less fat mass and more lean mass compared to flox/flox controls after 10 weeks of IF (Figure S3). Specifically, adipoIrf4 mice lost less gWAT and iWAT mass and adipoIrf4 mice had smaller TA muscles compared to flox/flox mice after IF (Figure 4E-G). We also found that 8 weeks of IF lowered fasting insulin, HOMA-IR, and blood glucose during a glucose tolerance test (GTT) in flox/flox and adipoIrf4 mice (Figure S3). Overall, the data show that deletion of adipocyte *Irf4* decreased fat loss and increased muscle loss during IF in obese mice. However, IF can improve blood glucose control independent of adipocyte *Irf4*.

**Figure 4:**
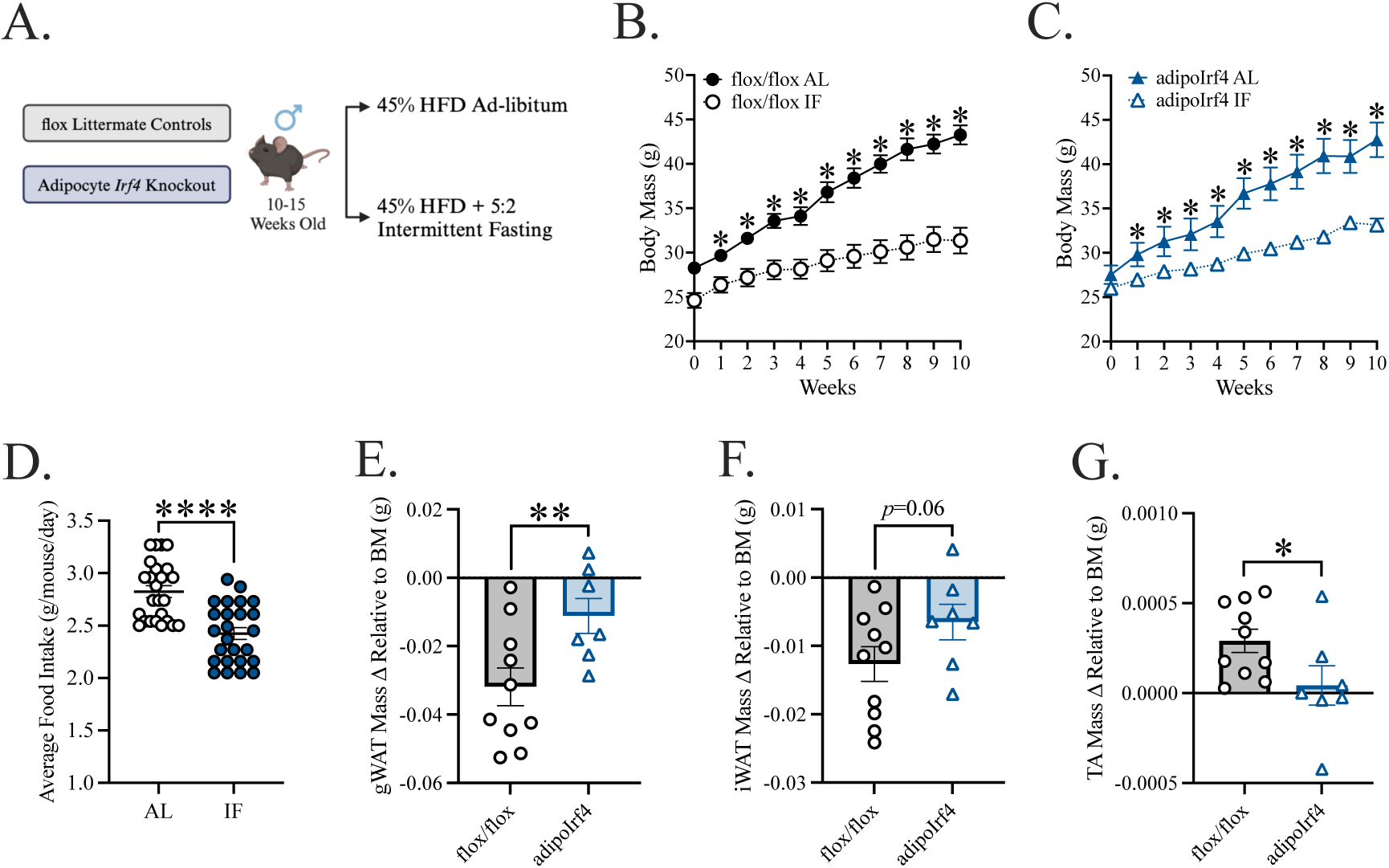
Adipocyte *Irf4* Impairs Lipolysis and Fat Loss While Promoting Muscle Loss in a Model of Reduced Food Intake During Intermittent Fasting. All HFD repeated IF experiments were also done at room temperature (23°C). flox/flox: *Irf4*^flox/flox^ control (*Irf4*^+/+^), adipoIrf4: adipocyte-specific knockout (*Irf4*^-/-^), and AL: ad-libitum, IF: intermittent fasting. Tissue weights were made relative to body mass and the fasting values were subtracted from the ad-libitum average of the corresponding group. A) 10wk 5:2 repeated intermittent fasting protocol. B) flox/flox AL and IF body mass. C) AdipoIrf4 AL and IF body mass. D) AL and IF average daily food intake. E) gWAT mass delta. F) iWAT mass delta. G) TA mass delta. Data presented as mean ± SEM (n=7-10). Statistical significance was determined using t-test analysis (A-F). *p≤0.05, **p≤0.01, and ****p≤0.0001.

### Deletion of adipocyte Irf4 decreased fat loss and increased muscle loss in an obese mouse model of IF with equal caloric intake

Next, we placed adipocyte-specific *Irf4* knockout mice (adipoIrf4) and littermate floxed mice (flox/flox) on an obesogenic 45% HFD, where mice were fed AL or underwent 5:2 IF for 10 weeks and housed at thermoneutrality (29°C), which was a model of equal food intake during IF (Figure 5A). IF decreased body mass in both adipoIrf4 and flox/flox mice (Figure 5B, C). However, average daily food intake was not different in AL and IF mice (Figure 5D). IF mice had increased activity (Figure 5E). AdipoIrf4 mice lost less gWAT mass and had smaller TA muscles compared to flox/flox mice after 10 weeks of IF (Figure 5F-H). AdipoIRF4 mice had larger adipocyte cross-sectional area and small TA muscle myofiber area in both AL and IF conditions (Figure 5I-N). Compared to flox/flox mice, AdipoIrf4 mice had no change in energy expenditure (Figure 5O). After IF, AdipoIrf4 mice had a higher respiratory exchange ratio (RER) in the fed state (Figure 5P). Hence, AdipoIrf4 mice had lower fat oxidation and increased carbohydrate oxidation after IF (Figure S4). 10 weeks of IF reduced liver fat and liver triglycerides in all mice (Figure S4). Overall, the data show that deletion of adipocyte *Irf4* decreased fat loss and increased muscle loss during an IF protocol that does not reduce food intake in obese mice. We found that adipoIrf4 mice had lower serum NEFA compared to flox/flox mice during IF (Figure 5Q).

**Figure 5:**
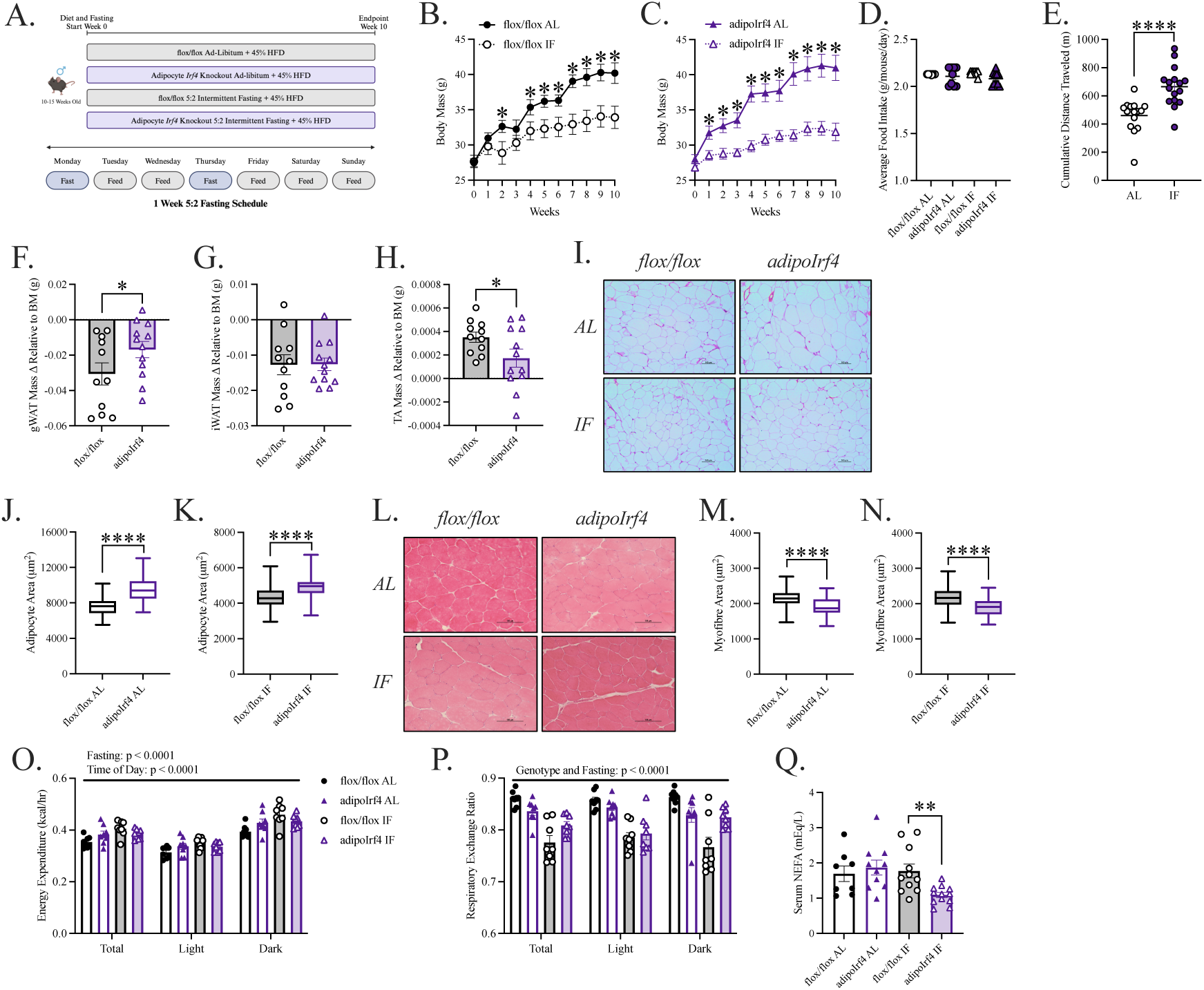
Adipocyte *Irf4* Impairs Lipolysis, Fat Loss and Oxidation, and Drives Muscle Loss in a Model of Equal Food Intake During Intermittent Fasting. All acute fasting and adrenergic experiments were done at room temperature (23°C) and mice were fed a chow diet. All HFD repeated IF experiments were done at thermoneutrality (29°C). flox/flox: *Irf4*^flox/flox^ control (*Irf4*^+/+^), adipoIrf4: adipocyte-specific knockout (*Irf4*^-/-^), and AL: ad-libitum, IF: intermittent fasting. Tissue weights were made relative to body mass and the fasting values were subtracted from the ad-libitum average of the corresponding group. A) 10wk 5:2 repeated intermittent fasting protocol. B) flox/flox AL and IF body mass. C) AdipoIrf4 AL and IF body mass. D) Average daily food intake. E) Cage activity over 24hrs. F) gWAT mass delta. G) iWAT mass delta. H) TA mass delta. I) Adipocyte images taken at 10X objective where the scale bar represents 100mm. J) Adipocyte cross-sectional area for AL mice. K) Adipocyte cross-sectional area for IF mice. L) Myofibre images taken at 10X objective where the scale bar represents 100mm. M) Myofibre cross-sectional area for AL mice. N) Myofibre cross-sectional area for IF mice. O) Average energy expenditure over 24hrs of feeding. P) Average respiratory exchange ratio over 24hrs of feeding. Q) Fed serum NEFA. Data presented as mean ± SEM (n=10-12). Statistical significance was determined using t-test analysis (B-N and Q). Significance was determined with an ANCOVA (O) and two-way ANOVA (P). *p≤0.05, **p≤0.01, and ****p≤0.0001.

## Discussion

IF has become popular because of purported metabolic health benefits (26). IF may decrease body mass in some people but, it is important to understand the relative proportion of muscle or adipose tissue that is lost. People with obesity may try IF, but it appears a feature of obesity may tip the balance toward muscle loss versus fat loss during IF. We recently showed that IF increases fat oxidation in lean mice but not in a mouse model of obesity and T2D (27). It is not clear how features of obesity/T2D or insulin resistance alter substrate use during fasting, but this is important because they may dictate the balance of fat mass loss versus muscle mass loss during IF. As fasting proceeds, the relative contribution of substrates progresses from liver glycogen to lipids/ketones, and some gluconeogenic amino acids (26). We found that humans with obesity and higher blood insulin had changes in blood levels of amino acids, including lower serum alanine (a key gluconeogenic substrate), and lost more lean mass compared to people without obesity during a 48 fast. We then used mice to manipulate insulin and determine how this altered muscle fat balance during IF. In mice, we used a 5:2 IF model over multiple weeks, to explore the longer-term impacts on relative at and muscle loss (28).

We hypothesized that higher insulin would suppress lipolysis, thus lowering lipids and glycerol as fuel sources to maintain glycemia during fasting, thereby lowering adipose loss during IF. We found that insulin-responsive adipocyte IRF4 was a key factor regulating the balance of skeletal muscle versus fat loss during IF in mice. It was already known that adipocyte-specific deletion of *Irf4* equates to less fat mass loss during an acute fast in mice (22). IRF4 can regulate lipolysis through adipose triglyceride lipase (ATGL) and hormone sensitive lipase (HSL) expression and adipocyte-specific deletion of *Irf4* lowers serum NEFA and glycerol after stimulation with the b3-agonist isoproterenol (22). Insulin can regulate lipolysis transcriptionally via cytosolic sequestration of *FoxO1* and subsequent reduction of *Irf4* expression, which reduces *Atgl* and *Hsl* expression in adipose tissue (22, 23). Hence the role of insulin on multiple branches of lipolysis should be considered in the regulation of adipose loss (versus muscle loss) during IF. In addition, the impact of insulin and IRF4 on other processes beyond lipolysis should be considered in regulating fat and muscle mass (i.e., ketogenesis or proteolysis). IRF4 can regulate muscle glycogen and glucose metabolism in muscle (29). In addition, there is metabolic communication between muscle and liver that is regulated by IRF4 (30). Hence, adipose tissue IRF4 communication with muscle (and the liver) during IF should also be investigated.

A key finding from human data was lean mass loss after 48 hours of acute fasting. Individuals with obesity lost more total body mass and lean mass compared to lean controls. Recent evidence in humans undergoing a 7-day fast show that lean individuals with a BMI of 25kg/^2^ lose ∼2kg of body mass after 48 hours of the protocol, which aligns with our data. Furthermore, after 7-days of fasting lean individuals can lose up to 6kg in total body mass, with 4kg from lean mass and 2kg from fat mass (31). This data provides a foundation for the idea that obesity and/or high insulin drives lean mass loss during fasting and aligns with findings from our hyperinsulinemic mouse model and chronic IF. Additionally, a single fast in humans showed that individuals living with obesity had lower serum alanine levels, which may indicate increased activity of the glucose-alanine (Cahill) cycle (32, 33). The glucose-alanine cycle is upregulated in obesity and T2D, where alanine is mobilized from muscle via ALT to fuel gluconeogenesis (34). This loss of amino acid from muscle may contribute to lower lean mass during fasting, which may compound during repeated IF. Future work should determine if insulin regulation of IRF4 alters the glucose-alanine cycle. Our results in mice do not show any differences in static serum levels of amino acid levels or alanine, but metabolic flux studies are warranted. Glucagon is known to promote proteolysis and liberate amino acids from muscle to fuel gluconeogenesis (35, 36). Importantly, we showed that glucagon was unchanged (or even lower) in mice with hyperinsulinemia that lost more muscle and less fat during IF.

IRF4 regulates the thermogenic program in brown adipose tissue (BAT) (37). Both cold stimulation and cAMP increased IRF4 in murine and human BAT. IRF4 directly interacts with peroxisome proliferator-activated receptor gamma coactivator 1 a (PGC1a) to increase uncoupling protein 1 (UCP1) expression in mice thereby increasing energy expenditure (37). Mice lacking *Irf4* in BAT have increased skeletal muscle myopathy and increased serum myostatin, a negative regulator of muscle growth (38). Restoring BAT *Irf4* decreased serum myostatin (38). These data suggest interorgan communication between adipose tissue depots and skeletal muscle with hormones such as myostatin can dictate fat loss versus muscle loss. It is enticing to speculate that insulin control of IRF4 in adipose tissue may dictate changes in muscle growth factors. To date, studies have not connected WAT IRF4 expression and myostatin, only BAT.

IF restricts the timing of food intake, but this does not always result in lower food intake over a longer term. In mice that undergo ADF or 5:2 IF there is hyperphagia during re-feeding days post-IF which equilibrates the weekly caloric intake (10, 39). This response does not always occur in humans (8). Various forms of IF have been shown to increase energy expenditure through BAT activation or browning of WAT through various mechanisms independently of changes in food intake in mice (39–41). However, it is still unclear how repeated IF affects energy expenditure in humans. Importantly, we used two different models of IF and our results show that deletion of IRF4 within adipocytes promotes muscle loss over fat loss in mouse models of reduced caloric intake and equal caloric intake during IF. It is not clear why housing HFD-fed mice at thermoneutrality versus room temperature caused changes in food intake during 5:2 IF, but this finding warrants further study.

In summary, we show that features of obesity such as high insulin influence muscle versus fat balance during IF. Elevated systemic blood insulin led to less fat loss during IF. IRF4 was identified as an insulin-responsive factor that, when deleted in the adipocytes of mice, promoted more skeletal muscle loss and less fat loss during IF. Elevated insulin could be an indicator of increased risk for exacerbated muscle loss and less fat loss during IF in individuals with obesity. Given that obesity is linked to hyperinsulinemia, we propose that blood insulin levels should be measured and carefully considered in the clinic before and during IF, if individuals with obesity pursue or are prescribed IF for weight loss.

## Materials and Methods

### Human Fasting Experimental Model

The present paper reports secondary outcomes from a two-group pre-post experimental trial in participants with lean BMI or living with obesity. This trial was approved by the University of British Columbia Clinical Research Ethics Board (H22-03605) and was registered with ClinicalTrials.gov (NCT05886738). All participants provided written informed consent prior to participating this trial.

### Participants

Participants were included if they had a BMI between 18.5 and 24.9 kg/m^2^ (group with lean BMI) or 30 kg/m2 (group living with obesity), were ages 19-69 years, accumulated less than 150 minutes of moderate-to-vigorous physical activity per week, and were able to read and understand English to follow the study procedures and provide written informed consent. Participants were excluded if they had an autoimmune or inflammatory disease diagnosis, a cancer diagnosis and/or treatment within the five years prior, diagnosed type 1 or 2 diabetes, a history of cardiovascular events, or pregnant. Participants were excluded if they were taking glucose-lowering medications, thyroid medications, or any other medications known to affect glucose or lipid metabolism. Additionally, participants were excluded if they currently smoked cigarettes or were unwilling to refrain from using cannabis for the duration of the study. Finally, individuals regularly practicing intermittent fasting, currently following a ketogenic diet, or actively losing/gaining weight (defined as >4 kg weight loss/gain in the last month) were excluded from the study.

### Study Visit Procedures

Participants arrived having fasted overnight (∼10 hours) and avoided exercise, alcohol, and caffeine the day prior. After the study procedures were reviewed and written consent had been obtained, anthropometric data was collected, and body composition was analyzed using the Tanita TBF-410GS body composition analyzer. Participants consumed a standardized meal replacement drink (Ensure Plus Calories) and a baseline fed venous blood sample was collected 2 hours later. After participants had fasted for 48 hours, they returned to the lab for their final visit, during which anthropometrics, body composition, and a final venous blood sample were obtained.

### Blood Sampling

Serum was collected into Serum Separator Tubes (BD Vacutainer) and allowed to clot for 1 hour, after which it was centrifuged for 15 minutes at 4°C and 2,000 g, aliquoted, and immediately frozen at -70°C for future batch analyses. Serum insulin was quantified in duplicate by ELISA according to the manufacturer’s directions. To assess lipolysis, serum NEFA was measured using commercially available kits. To assess proteolysis and amino acid catabolism, total serum L-amino acids and alanine were measured. To ensure proper detection of metabolites, samples were diluted 2X for NEFA, 2X for L-amino acids, and 2X for alanine.

### Murine Models and Genotyping

All animal procedures for this study were approved by the McMaster University Animal Research Ethics Board in accordance with the guidelines of the Canadian Council of Animal Care. All mice were 10-15 weeks old before experiment initiation. Mice were maintained on a 12-hour light/dark cycle in a specific-pathogen-free (SPF) facility, and experiments were performed on multiple cohorts of mice born from different parents across different times of the year. Wild-type C57BL/6J, Adipoq-Cre^+^, *Irf4*^flox^/J, and whole-body *Irf4*^KO^/J mouse strains were obtained from Jackson Laboratories and bred in-house (heterozygous x heterozygous) to produce littermate control and knockout animals. When breeding animals with the Cre recombinase, only one parent, male or female, contained the Adipoq-Cre promotor to ensure litter viability and consistency.

*Irf4* deletion was confirmed by tail DNA genotyping. All primers were made in-house at the Mobix Lab in the McMaster Genomic Facility. Mouse tail clippings (2-3 mm) were digested using a DNA Fast Extract kit (Wisent Advanced, 801-200-HR) according to manufacturer’s instructions. Presence of the Adipoq-Cre transgene was confirmed by PCR amplification of isolated DNA. Primers for the Adipoq-Cre and a PCR internal control amplification band were used in a single reaction. In a separate reaction, the presence of *Irf4^flox^* was confirmed by PCR amplification. Presence or absence of these genes was validated using gel electrophoresis and a 1% agarose gel. Detection of the *Irf4* gene in our whole-body knockout model was validated using specific primers for *Irf4*-intact, *Irf4*-heterozygous, and *Irf4*-knockout sequences. These genes were validated using gel electrophoresis and a 2% agarose gel run in 1% TAE buffer at 100V for 3 hours. For the *Irf4* whole-body KO model, the wild-type band is detected at 150bp and the KO band at 500bp. For the adipocyte *Irf4* KO model, *Irf4*flox was detected at 91bp. For adipocyte *Irf4* models, flox animals were crossed with Cre+ animals, and the Adipoq-Cre transgene was detected at 200bp, with an internal positive control band detected at 324bp. Tissue specific KO animals were only deemed full KO and used if both alleles possessed flox sites in cells that were present for the Cre recombinase (42).

### High Chronic Insulin Model

To model chronic increased peripheral insulin, two LinBit insulin pellets were surgically implanted in the backs of mice, which chronically release a set amount of insulin per day at a dose of 0.1 U/day/pellet. Mice were first anesthetized using isoflurane before having the mid-back region shaved and sanitized with 10% iodine solution. After the iodine was applied, it was wiped off using isopropyl alcohol and dried with sterile gauze. Insulin pellets were then subcutaneously inserted into the mid-back region of the mice using a trocar provided by the manufacturer, and the insertion site was sealed using tissue glue provided by the McMaster Central Animal Facility.

Following the surgical procedure, mice were individually housed to avoid aggravation of the implant site for both acute and chronic fasting experiments. For acute fasting studies, the insulin pellets were left in the mice 7 days prior to the start of the experiments, and for long-term intermittent fasting experiments, pellets were left in the mice 3 days prior. This was done as the pellets last for ∼40-42 days or 6 weeks before needing to be changed to ensure their potency. For long-term intermittent fasting studies, the pellets were changed during week 6 of the 10-week fasting protocol. Blood glucose was measured weekly to ensure sufficient insulin levels throughout the study.

### Intermittent Fasting Protocol

Prior to the fasting protocol, mice were housed at room temperature (23°C) in groups of up to 5 mice per cage and fed a standard chow diet (Teklad 22/5 Rodent Diet). At the initiation of the 5:2 intermittent fasting protocol, mice were switched to a 45% kcal from fat diet (Research Diets D12451) and either kept at room temperature (23°C) or moved into Solace Zone (ARES Scientific) heated cages, set to 29°C, to simulate thermoneutral conditions.

5:2 intermittent fasting was performed on Mondays and Thursdays for a total of 10 weeks. Mice either had ad-libitum food access or total food deprivation for 24 hours, and all groups had full water access. Intermittent fasting mice were food deprived from 11:00am to 11:00am of the following day to implement periods of 24-hour fasting and feeding. During the fast, bedding was changed for both intermittent fasting and ad-libitum mice to fully eliminate food residue and avoid variable treatment of the groups. Weekly food intake, blood glucose, and body weight measurements were taken on Monday before the first fast of the week. To measure daily food intake, we monitored the weight of the remaining food in the cage every Monday and averaged by mouse number. Blood glucose was monitored via tail vein blood (StarStrip Xpress glucometer and strips). Body composition measures were acquired using time-domain nuclear magnetic resonance (TD-NMR) (Bruker Minispec Live Mice Analyzer) on Monday of week 0 and week 8 of the protocol to gauge body, lean, and fat mass. Metabolic cage measures (Sable Systems) were taken on week 9 of the study. Animals were left to acclimatize to the thermoneutral metabolic cages for 2-3 days before taking fed and fasted measurements. Fed and fasted serum was collected on Wednesday and Thursday of week 10 using tail vein blood that was collected in heparin-lined capillary tubes and stored on ice before being centrifuged at 10, 000g for 10 minutes at 4°C.

### In Vivo Injection Tests

Mice were fed a standard chow diet and the lipolytic test was done in the fed state at 9:00am. Acute lipolysis was stimulated using intraperitoneal injection of 0.033nmol/g CL-316, 243 (Tocris Biosciences). This lower dose of CL-316, 243 has been shown to promote a lipolytic response through activation of b3-adrenergic receptors without promoting excessive inflammation, which can also drive lipolysis (43–46). Blood was taken at 0, 20, and 60 minutes following the injection, stored on ice, and centrifuged at 10, 000g for 10 minutes at 4°C to obtain serum.

On Thursday of week 8 of the *Irf4* KO IF trials, glucose tolerance of mice was measured using an intraperitoneal glucose tolerance test (GTT). Mice were subject to a 6-hour fast, which replaced the 24-hour fast for that week, from 8:00am to 2:00pm, followed by a fasting blood glucose measurement. Mice were then injected intraperitoneally with a 1.5g/kg solution of D-glucose, and blood glucose was measured at 0, 20, 30, 40, 60, 90, and 120 minutes. Analysis of blood glucose levels at each time point, as well as the overall area under the curve, was completed for each group.

### Blood Assays

To assess lipolysis, serum NEFA and glycerol were measured using commercially available kits. To assess proteolysis and amino acid catabolism, total serum L-amino acids and alanine were measured. To ensure proper detection of metabolites, samples were diluted 4X for NEFA, 2X for glycerol, 10X for L-amino acids, and 2X for alanine. Fed and fasted serum samples were diluted 80X with assay buffer to fit within the provided standard curve. All assays were quantified using a Synergy H4 Hybrid reader (Biotek).

### RT-qPCR

RNA was extracted from ∼50-100mg of tissue for gene expression analysis. Ceramic beads were added to the tubes to mechanically homogenize the tissue at 5m/s for 60 seconds using a FastPrep-24 Tissue Homogenizer (MP Biomedicals). Once the tissue was homogenous, several steps were used to isolate pure RNA, including chloroform-phenol extraction, isopropanol precipitation, ethanol cleaning, dissolving the RNA pellet in ultra-pure water, and quantitation via microplate reader detection (BioTek). RNA concentrations across samples were equalized such that 500ng-1000ng total RNA was used for cDNA preparation. RNA was treated with DNase I, and cDNA was synthesized with SuperScript IV Reverse Transcriptase. Expression of transcripts for each gene was measured with TaqMan Gene Expression Assays using AmpliTaq Gold DNA Polymerase. Target gene expression was compared to the housekeeping gene expression of *Rplp0* using the ^ΔΔ^CT method.

### Liver and Muscle Assays

For all liver assays, ∼50mg of frozen liver was homogenized in 200mL of each respective assay buffer and ceramic beads using a FastPrep-24 Tissue Homogenizer (MP Biomedicals) set to 5m/s for 60 seconds. Liver triglycerides were assessed using a commercially available kit following the manufacturer’s protocol. Liver samples were diluted 50X with assay buffer to fit within the specified standard curve. Liver glycogen was assessed using a commercially available kit following the manufacturer’s protocol. Liver samples were diluted 50X with assay buffer to fit within the specified standard curve. Liver ALT activity was also assessed with a commercially available kit where the manufacturer’s protocol was followed. Several dilutions were tested, and liver samples were diluted 10X with assay buffer to fit within the specified standard curve.

Muscle ALT activity was assessed using a commercially available kit where the manufacturer’s instructions were followed. Samples were prepared using ∼25mg of TA muscle and homogenized in 200mL ALT assay buffer using ceramic beads and a tissue homogenizer set to 5m/s for 60 seconds. After testing dilutions, TA samples were diluted 2X to provide the best signal. All assays were quantified using a microplate reader (Biotek).

### Histological Analysis

Muscle samples were excised, weighed, dipped in tissue freezing medium, and immediately hardened in liquid nitrogen-cooled isopentane before being snap-frozen in liquid nitrogen and stored at -80°C. Liver and adipose sections were also excised, weighed, and immediately preserved in 10% formalin solution with 90% PBS for 72 hours before being transferred to 70% ethanol and stored at 4°C. 10μm muscle sections of the tibialis anterior muscle were cut with the Leica CM1850 Cryostat, maintained at -20°C, before being stained with H & E. Liver and adipose samples were sent to the John Mayberry Histology Facility at McMaster University to be paraffin-embedded, sectioned into 5μm slices, and stained with H & E. Images of liver, adipose, and muscle were acquired using a Nikon Eclipse Ti microscope, DS-Qi2 camera, and images were analyzed using Nikon NIS-Elements software. Researchers were blinded to the experiment conditions during image analysis. 100 cells were circled per image/mouse, with 25 adjacent cells from 4 different areas of the image used to gain a representative median cell size. Adipocyte and myocyte cell areas were calculated and binned relative frequencies of cell sizes were graphed.

### Statistical Analysis

We anticipated a medium-to-large effect size for the primary comparison between lean and obesity for body composition and serum metabolite changes in response to fasting in humans. Using an effect size of d=1.0, a two-tailed alpha of 0.05 and 80% power, N=17 per group was estimated for an independent sample t-test using G*Power v3.1. To obtain balanced numbers of males and females in each group, N=16 per group was recruited with 8 males and 8 females per group to underscore any potential sex differences. Statistical significance between two groups was determined using unpaired Welch’s t-tests. Where 3 or more groups were analyzed, a one-way or two-way ANOVA test was used with Tukey’s multiple comparisons, depending on the number of independent variables. In the case of time course experiments with two groups, multiple t-test analysis was used with a Welch correction. To test for normality, a Shapiro-Wilk test was conducted. Statistical analyses and sample size (n) for each experiment and graph are reported in the figure legends. If outliers were detected via a ROUT test, they were removed from the data set. Data are expressed as means ± SEM with significance set at p ≤ 0.05. Data was analyzed and graphs were made using GraphPad Prism 9 statistical software.

## Acknowledgments

This work was supported by a grant from the Natural Science and Engineering Research Council (NSERC, RGPIN-2020-05707) to Jonathan D. Schertzer. Jonathan D. Schertzer holds a Canada Research Chair in Metabolic Inflammation. Daniel M. Marko was supported by CIHR CGS-D and Michael DeGroote PhD awards.

## Supplementary Information

Supplemental information includes resource/reagent table and 4 supplemental figures. Specifically, S1-S4 contains data underlying the display items in the manuscript, related to Figures 2-5.

