## Supplemental Information for "Insulin Prevents Fat Loss and Promotes Muscle Loss During Intermittent Fasting in Obesity"

**Supplemental Materials**

**Supplemental Figure Legends: S1-S4**

**Supplementary Materials**

| REAGENT or RESOURCE | SOURCE | IDENTIFIER |
|---|---|---|
| **Chemicals, Peptides, and Recombinant Proteins** | | |
| CL-316, 243 | Tocris Bioscience | Cat# 1499 |
| D-Glucose | Sigma-Aldrich | Cat# G7021 |
| LinBit Insulin Pellets | LinShin | Cat# Re-1-T |
| Alanine | Sigma-Aldrich | Cat# A7627 |
| TRIzol | Invitrogen | Cat# 15596018 |
| Buffered Formalin (10%) | ACP Chemicals | Cat# F6000 |
| Hematoxylin Stain | ThermoFisher Scientific | Cat# GHS232 |
| Eosin Y Stain | ThermoFisher Scientific | Cat# 318906 |
| Xylene | ThermoFisher Scientific | Cat# 53056 |
| **Critical Commercial Assays** | | |
| NEFA-HR(2) Assay | Fujifilm Wako Chemicals | Cat# 434-91795, 436-91995, 270-77000 |
| Glycerol Assay | Sigma-Aldrich | Cat# F6428 |
| Murine Insulin ELISA | Toronto Bioscience | Cat# 32270 |
| Triglyceride Assay | Abcam | Cat# ab65336 |
| Glycogen Assay | Abcam | Cat# ab169558 |
| L-Amino Acids | Sigma-Aldrich | Cat# MAK002 |
| Alanine | Sigma-Aldrich | Cat# MAK001 |
| ALT Assay | Sigma-Aldrich | Cat# MAK052 |
| β-hydroxybutyrate | Sigma-Aldrich | Cat# MAK041 |
| Glucagon ELISA | Mercodia | Cat# 10-1281-01 |
| Human Insulin ELISA | Cayman Chemical | Cat# 90095 |
| **Experimental models: Organisms/Strains** | | |
| C57BL/6J | Jackson Laboratory | Cat# 000664 |
| B6.129P2-*Irf4*$^{tm1Mak}$/J (*Irf4*$^{KO}$*/J*) | Jackson Laboratory | Cat# 031834 |
| B6.129S1-*Irf4*$^{tm1Rdf}$/J (*Irf4*$^{flox}$*/J*) | Jackson Laboratory | Cat# 009380 |
| B6.FVB-Tg(Adipoq-cre)1Evdr/J (*Adipoq-Cre*$^{+}$*/J*) | Jackson Laboratory | Cat# 028020 |
| **Oligonucleotides** | | |
| B6.129P2-*Irf4*$^{tm1Mak}$/J (*Irf4*$^{KO}$*/J*):<br>WT1 Seq: GCA ATG GGA AAC TCC GAC AGT<br>WT2 Seq: CAG CGT CCT CCT CAC GAT TGT<br>MUT1 Seq: CCG GTG CCC TGA ATG AAC TGC<br>MUT2 Seq: CAA TAT CAC GGG TAG CCA ACG | Mobix Lab McMaster Genomics Facility | N/A |
| B6.129S1-*Irf4*$^{tm1Rdf}$/J (*Irf4*$^{flox}$*/J*):<br>Flox1 Seq: TGG GCA CCT CTA CTG TCT GG<br>Flox2 Seq: CTC TGG GGA CAT CAG TCC T<br>Flox3 Seq: CGA CCT GCA GCC AAT AAG C | Mobix Lab McMaster Genomics Facility | N/A |

| | | |
|---|---|---|
| B6.FVB-Tg(Adipoq-cre)1Evdr/J (*Adipoq-Cre*$^{+}$*/J*):<br>Cre1 Seq: ACG GAC AGA AGC ATT TTC CA<br>Cre2 Seq: GGA TGT GCC ATG TGA GTC TG<br>Cre3 Seq: CTA GGC CAC AGA ATT GAA AGA TCT<br>Cre4 Seq: GTA GGT GGA AAT TCT AGC ATC ATC C | Mobix Lab McMaster Genomics Facility | N/A |
| *Irf4* Taqman Mouse Primers | ThermoFisher Scientific | Cat# Mm00516431 |
| *Rplp0* Taqman Mouse Primers | ThermoFisher Scientific | Cat# Mm01974474 |
| **Software and Algorithms** | | |
| ImageJ 1.46r | NIH | RRID:SCR_003070 |
| NIS-Elements | Nikon | RRID:SCR_002776 |
| GraphPad Prism Software, Version 9/10 | GraphPad Software | RRID:SCR_002798 |
| **Other** | | |
| Standard Chow Diet | Envigo | Cat# 8640 |
| 45% kcal from fat Diet (HFD) | Research Diets | Cat# D12451 |
| Heparinized Micro-Hematocrit Capillary Tubes | Fisher Scientific | Cat# 22-362-566 |
| Histology Cassettes | Simport Scientific | Cat# M498-6 |
| Tissue Freezing Medium (OCT) | General Data Healthcare | Cat# TFM-C |
| 2x HS-Red Taq Mix | Wisent | Cat# 801-200-HR |
| DNA Fast Extract Buffer A | Wisent | Cat# 801-200-HR |
| DNA Fast Extract Buffer B | Wisent | Cat# 801-200-HR |
| DNase I | Invitrogen | Cat# 18068015 |
| Superscript IV | Invitrogen | Cat# 18090010 |
| AmpliTaq Gold DNA Polymerase, Gold Buffer, and $MgCl_2$ | Applied Biosystems | Cat# 4311820 |

### Supplementary Figures

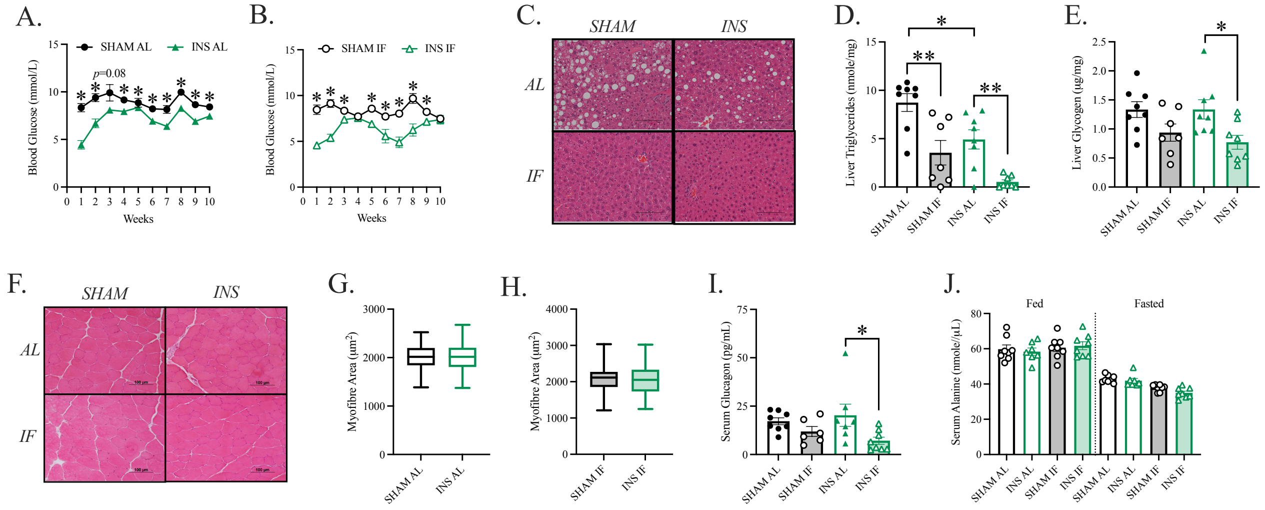


**Supplemental Figure 1: Supplemental Data in Support of Figure 2.** SHAM: control, INS: hyperinsulinemia, AL: ad-libitum, IF: intermittent fasting. Delta values from MRI were calculated where intermittent fasting values were subtracted from the ad-libitum average of the corresponding group. A) Weekly fed blood glucose in AL mice. B) Weekly fed blood glucose in IF mice. C) Hepatocyte images taken at 10X objective where the scale bar represents 100µm. D) 24hr fasted liver triglycerides. E) 24hr fasted liver glycogen. F) Myofiber images taken at 10X objective where the scale bar represents 100µm. G) Myofibre cross-sectional area for AL mice. H) Myofibre cross-sectional area for IF mice. I) 24hr fasted serum glucagon. J) Fed and 24hr fasted serum alanine. Statistical significance was determined using t-test analysis (A-F and K, L). Significance was determined using a one-way ANOVA (H, I, M). *$p \leq 0.05$, **$p \leq 0.01$, and ***$p \leq 0.001$.

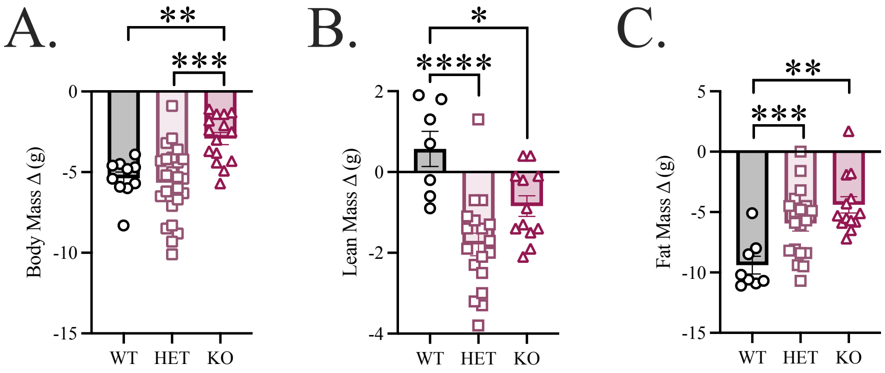


**Supplemental Figure 2: Supplemental Data in Support of Figure 3.** WT: wild-type control, HET: heterozygous (*Irf4*$^{+/-}$), KO: *Irf4* knockout (*Irf4*$^{-/-}$), and AL: ad-libitum, IF: intermittent fasting. Delta values from MRI were calculated where intermittent fasting values were subtracted from the ad-libitum average of the corresponding group. A) WT, HET, and KO body mass delta. B) WT, HET, and KO lean mass delta. C) WT, HET, and KO fat mass delta. Data presented as mean ± SEM (n=10-25). Statistical significance was determined using a one-way ANOVA (A-C). *p≤0.05, **p≤0.01, ***p≤0.001, and ****p≤0.0001.

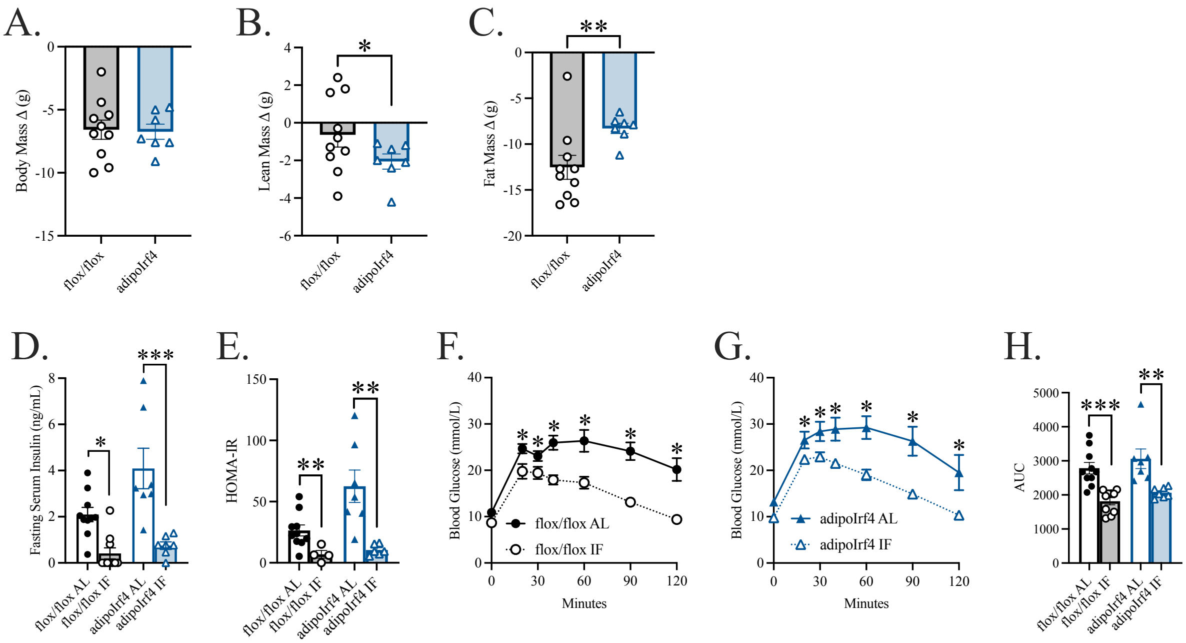


**Supplemental Figure 3: Supplemental Data in Support of Figure 4.** flox/flox: *Irf4*$^{flox/flox}$ control (*Irf4*$^{+/+}$), adipoIrf4: adipocyte-specific knockout (*Irf4*$^{-/-}$), and AL: ad-libitum, IF: intermittent fasting. Delta values from MRI were calculated where intermittent fasting values were subtracted from the ad-libitum average of the corresponding group. A) flox/flox and adipoIrf4 body mass delta. B) flox/flox and adipoIrf4 lean mass delta. C) flox/flox and adipoIrf4 fat mass delta. D) flox/flox and adipoIrf4 AL and IF 6hr fasted serum insulin. E) flox/flox and adipoIrf4 AL and IF HOMA-IR. F) flox/flox AL and IF GTT. G) AdipoIrf4 AL and IF GTT. H) flox/flox and adipoIrf4 AL and IF AUC. Data presented as mean ± SEM (n=7-10). Statistical significance was determined using t-test analysis (A-C, F-H). Significance was determined with a one-way ANOVA (D, E). *p≤0.05, **p≤0.01, and ***p≤0.001.

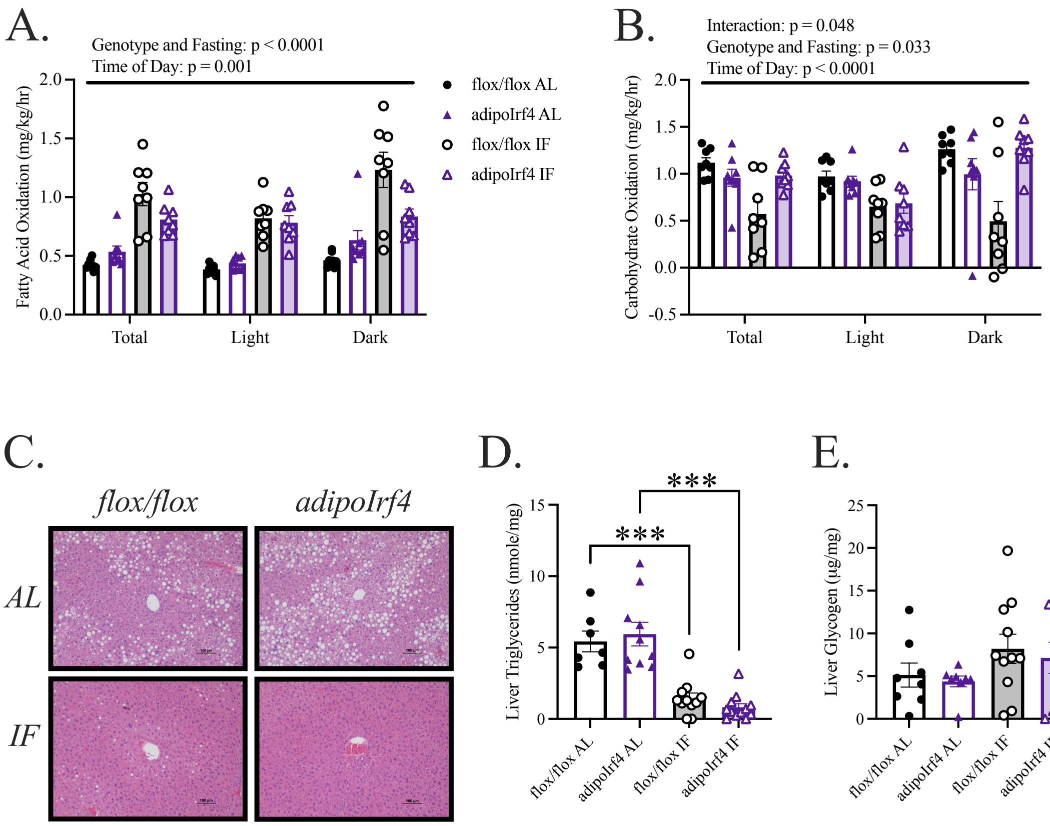


**Supplemental Figure 4: Supplemental Data in Support of Figure 5.** flox/flox: *Irf4*$^{flox/flox}$ control (*Irf4*$^{+/+}$), adipoIrf4: adipocyte-specific knockout (*Irf4*$^{-/-}$), and AL: ad-libitum, IF: intermittent fasting. Delta values from MRI were calculated where intermittent fasting values were subtracted from the ad-libitum average of the corresponding group. A) Average fatty acid oxidation over 24hrs of feeding. B) Average carbohydrate oxidation over 24hrs of feeding. C) Hepatocyte images taken at 10X objective where the scale bar represents 100µm. D) 24hr fasted liver triglycerides. E) 24hr fasted liver glycogen. Data presented as mean ± SEM (n=10-12). Statistical significance was determined with a two-way ANOVA (A, B) and one-way ANOVA (D, E). ***p≤0.001.
